# PoSeiDon: a Nextflow pipeline for the detection of evolutionary recombination events and positive selection

**DOI:** 10.1101/2020.05.18.102731

**Authors:** Martin Hölzer, Manja Marz

## Abstract

**Summary:** PoSeiDon is an easy-to-use pipeline that helps researchers to find recombination events and sites under positive selection in protein-coding sequences. By entering homologous sequences, PoSeiDon builds an alignment, estimates a best-fitting substitution model, and performs a recombination analysis followed by the construction of all corresponding phylogenies. Finally, significantly positive selected sites are detected according to different models for the full alignment and possible recombination fragments. The results of PoSeiDon are summarized in a user-friendly HTML page providing all intermediate results and the graphical representation of recombination events and positively selected sites.

**Availability and implementation:** PoSeiDon is freely available at https://github.com/hoelzer/poseidon. The pipeline is implemented in Nextflow with Docker support and processes the output of various tools.

## 1 INTRODUCTION

Selection pressure continuously influences the evolution of genes and can be studied in many ways (Vitti et al., 2013). For example, positive or diversifying selection can be detected by comparing the rates of non-synonymous (*dN*) and synonymous substitutions (*dS*) in an alignment of orthologous genes. Over several sites (codons), the *dN/dS* ratio (or *ω*) can reach values well above 1 (Yang, 2007), and such sites are likely to be positively selected. For instance, specific amino acid changes are favored if they increase the host’s fitness against a pathogen (Fumagalli et al., 2011). Alternatively, the genes of a pathogen are affected, as in the COVID 19 pandemic, where positively selected sites in the spike protein of the virus gave cause for concern (Korber et al., 2020). The detection of positive selection enables researchers to gain insights into the evolution of genes and thus develop countermeasures against pathogens that are in a constant ‘arms-race’ with their host.

Since recombination can have a profound influence on evolutionary processes and can adversely affect phylogenetic reconstruction and the accurate detection of positive selection (Shriner et al., 2003), screening for breakpoints to define recombinant parts within an alignment should be a standard step in any comparative evolutionary study.

A comprehensive evolutionary analysis of significantly positively selected sites consist of several complicated steps, including (1) in-frame alignment; (2) Indel correction; (3) phylogenetic tree calculation; (4) selection of a best-fitting nucleotide substitution model; (5) detection of topological incongruence and breakpoint selection to describe putative recombination events; (6) calculation of positively selected sites (*ω >* 1) under varying models; (7) and their impact on the selective pressure acting on the whole alignment. Thus, such an analysis involves dozens of different tools and parameter settings.

In addition, the results of many well-established and widely used tools in this field of evolutionary science are not easy to interpret and process. Especially, the accurate detection and handling of putative recombination events is a challenging but essential task.

Currently, only a few tools for the comprehensive detection of positive selection exist. These tools either do not automatically combine all of the described steps (Delport et al., 2010), do not take possible recombination events into account (Doron-Faigenboim et al., 2005; Stern et al., 2007; Webb et al., 2017), or focus only on the detection of positive selection in prokaryotic genomes (Su et al., 2013).

Here we present PoSeiDon, a pipeline that allows researchers to perform comprehensive evolutionary studies by automatically taking care of all tasks mentioned above. PoSeiDon does detect not only positively selected sites in an alignment of homologous sequences but also possible recombination events that could otherwise adversely affect the positive selection detection. The input is a single FASTA file consisting of protein-coding DNA sequences with a correct open reading frame. The output is summarized in TEX and PDF format and visualized in a user-friendly HTML page, providing access to all results and intermediate files.

## 2 PIPELINE AND IMPLEMENTATION

PoSeiDon comprises an assembly of different scripts and tools (Fig. 1) that allow for the detection of recombination and positive selection in protein-coding sequences. Each third-party tool encapsulates in a Docker container and all steps are connected in a Nextflow (Di Tommaso et al., 2017) implementation for full parallelization and simple execution. If Nextflow and Docker are configured, PoSeiDon can be installed and run with a single command: `nextflow run hoelzer/poseidon --help`. Different profiles allow the reliable execution of PoSeiDon in the cloud or on a cluster system. Starting from homologous coding sequences provided by the user, we build a multiple sequence alignment guided by amino acid information with TranslatorX (v1.1) (Abascal et al., 2010), using Muscle (v3.8.31) (Edgar, 2004) to align the sequences. The resulting in-frame nucleotide alignment is cleaned for Indels. A best-fitting substitution model is selected using MODELTEST (Posada and Crandall, 1998), which is part of the HyPhy suite (v2.2.7) (Pond et al., 2005).

**Fig. 1.**
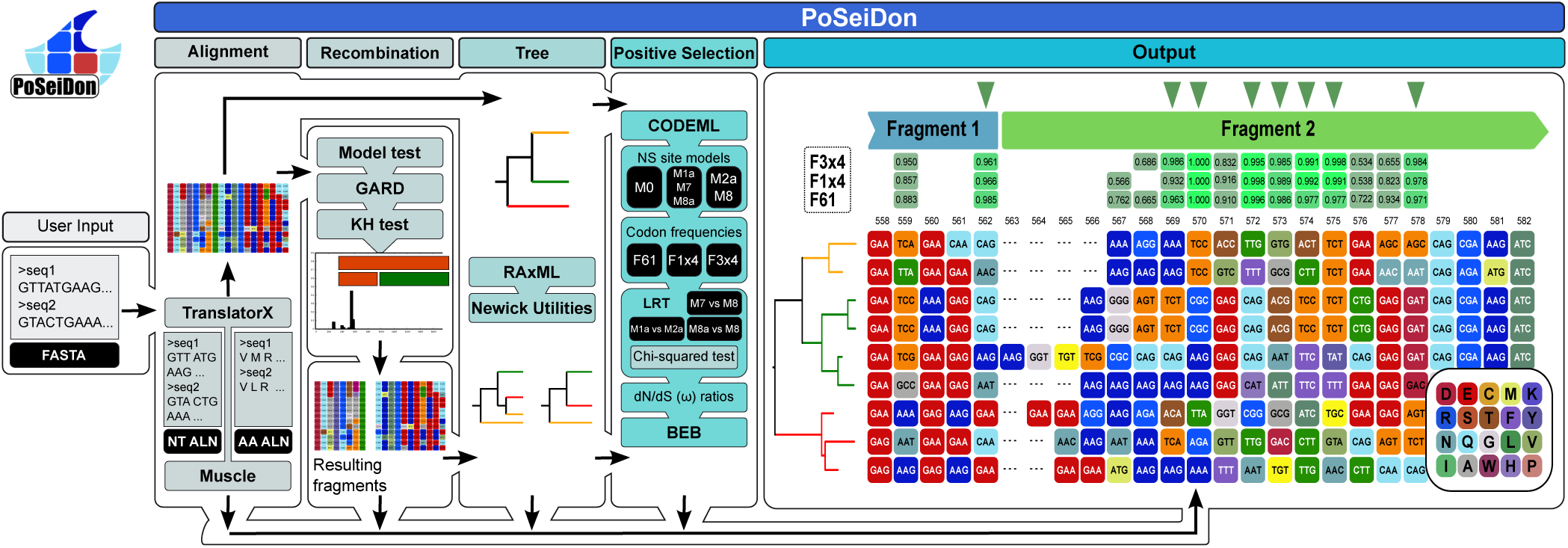
The workflow of the PoSeiDon pipeline and example output. The PoSeiDon pipeline comprises in-frame alignment of homologous protein-coding sequences, detection of putative recombination events and evolutionary breakpoints, phylogenetic reconstructions and detection of positively selected sites in the full alignment and all possible fragments. Finally, all results are combined and visualized in an HTML web page. The resulting alignment fragments are indicated with colored bars in the HTML output.

Possible recombination events and corresponding breakpoints in the alignment are detected using GARD (Pond et al., 2006b,a) under the previously selected substitution model. All breakpoints are tested for significant topological incongruence using a Kashino Hasegawa (KH) test (Kishino and Hasegawa, 1989). KH-insignificant breakpoints most frequently arise from variation in branch lengths between segments. However, we observed interesting positively selected sites in fragments without any significant topological incongruence (Fuchs et al., 2017). Thus, KH-insignificant breakpoints can be taken into account and are marked in the final output, as they might not occur from real recombination events. Positions of putative breakpoints that would destroy the open reading frame are adjusted to the next valid position. Phylogenetic reconstructions on the full alignment and all fragments are performed with RAxML (v8.2.12) (Stamatakis, 2014) using the `GTRGAMMA` model for nucleotide sequences and `PROTGAMMAWAG` for amino acids. All calculations are performed with 1 000 bootstrap replicates. The user can apply optional outgroup rooting. The Newick Utilities suite (v1.6) (Junier and Zdobnov, 2010) is used to visualize the calculated trees in different formats.

Positive selection is analyzed on the full alignment and each of the fragments separately. Maximum-likelihood tests to detect positive selection under varying site models are performed with CODEML (M1a vs. M2a, M7 vs. M8) implemented within the PAML suite (v4.9) (Yang, 2007). Furthermore, we implemented the M8a vs. M8 test proposed by Swanson et al. (2003) as an additional model test in PoSeiDon. The different statistical site models that do not allow (neutral models) or allow (selection models) a class of codons to evolve with *ω >* 1 are compared. Furthermore, varying codon frequency models are applied to simulate different nucleotide substitution rates. A gene is declared to be positively selected if the neutral model can be rejected in favor of the positive selection model based on a likelihood ratio test. Then, a Bayes empirical Bayes (BEB) approach (Yang et al., 2005) is applied to calculate posterior probabilities (*PP*) that a codon comes from the site class with *ω >* 1. Positively selected sites with an assigned *PP >* 0.95 are depicted as significant.

We graphically summarize all positively selected sites under varying frequency models in the output (Fig. 1). Thus, we allow the user to investigate sites that would be dismissed from the output when using a *PP* threshold. For example, such sites could be located in regulatory domains of the final protein, yielding a lower *PP* value due to insufficient species sampling (McBee et al., 2015). The final output of PoSeiDon is based on a heavily modified version of the TranslatorX HTML output. The amino acid color code is adapted from TranslatorX.

## 3 CONCLUSIONS

Here we present PoSeiDon, an easy-to-use Nextflow pipeline for the accurate detection of site-specific positive selection and recombination events in protein-coding sequences. The input is a multiple FASTA file of homologous coding sequences that is automatically transferred into a codon-based alignment. Since recombination can have a profound impact on the evolutionary history of sequences, we initially check the alignment for topological incongruence to define putative recombination breakpoints. The whole evolutionary analysis of PoSeiDon is performed independently for the full alignment and all possible fragments. PoSeiDon automatically calculates maximum likelihood-based phylogenetic trees for all alignments, estimates *ω* values at each site, and computes their impact on the positive selection. All identified sites and their significance values are projected onto the codon and amino acid alignment of the input sequences to allow visual identification of evolutionary hot-spots with high *ω* values. Additionally, publication-ready PDF and LATEX tables are provided, including all breakpoints and significantly positively selected sites. All results are summarized in a user-friendly and clear manner, allowing researchers to study positive selection.

## REFERENCES

Abascal, F., Zardoya, R., and Telford, M. J. (2010). TranslatorX: multiple alignment of nucleotide sequences guided by amino acid translations. Nucleic Acids Res, 38(Web Server issue), W7–13.

Delport, W., Poon, A. F. Y., Frost, S. D. W., and Kosakovsky Pond, S. L. (2010). Datamonkey 2010: a suite of phylogenetic analysis tools for evolutionary biology. Bioinformatics, 26(19), 2455–2457.

Di Tommaso, P., Chatzou, M., Floden, E. W., Barja, P. P., Palumbo, E., and Notredame, C. (2017). Nextflow enables reproducible computational workflows. Nature biotechnology, 35(4), 316–319.

Doron-Faigenboim, A., Stern, A., Mayrose, I., Bacharach, E., and Pupko, T. (2005). Selecton: a server for detecting evolutionary forces at a single amino-acid site. Bioinformatics, 21(9), 2101–2103.

Edgar, R. C. (2004). MUSCLE: multiple sequence alignment with high accuracy and high throughput. Nucleic Acids Res, 32(5), 1792–1797.

Fuchs, J., Hölzer, M., Schilling, M., Patzina, C., Schoen, A., Hoenen, T., Zimmer, G., Marz, M., Weber, F., Müller, M. A., et al. (2017). Evolution and antiviral specificities of interferon-induced Mx proteins of bats against Ebola, Influenza, and other RNA viruses. Journal of virology, 91(15), e00361–17.

Fumagalli, M., Sironi, M., Pozzoli, U., Ferrer-Admetlla, A., Ferrer-Admettla, A., Pattini, L., and Nielsen, R. (2011). Signatures of environmental genetic adaptation pinpoint pathogens as the main selective pressure through human evolution. PLoS Genet, 7(11), e1002355.

Junier, T. and Zdobnov, E. M. (2010). The Newick utilities: high-throughput phylogenetic tree processing in the UNIX shell. Bioinformatics, 26(13), 1669–1670.

Kishino, H. and Hasegawa, M. (1989). Evaluation of the maximum likelihood estimate of the evolutionary tree topologies from DNA sequence data, and the branching order in Hominoidea. J Mol Evol, 29(2), 170–179.

Korber, B., Fischer, W., Gnanakaran, S. G., Yoon, H., Theiler, J., Abfalterer, W., Foley, B., Giorgi, E. E., Bhattacharya, T., Parker, M. D., et al. (2020). Spike mutation pipeline reveals the emergence of a more transmissible form of SARS-CoV-2. bioRxiv.

McBee, R. M., Rozmiarek, S. A., Meyerson, N. R., Rowley, P. A., and Sawyer, S. L. (2015). The Effect of Species Representation on the Detection of Positive Selection in Primate Gene Data Sets. Mol Biol Evol, 32(4), 1091–1096.

Pond, S. L. K., Frost, S. D. W., and Muse, S. V. (2005). HyPhy: hypothesis testing using phylogenies. Bioinformatics, 21(5), 676–679.

Pond, S. L. K., Posada, D., Gravenor, M. B., Woelk, C. H., and Frost, S. D. W. (2006a). Automated phylogenetic detection of recombination using a genetic algorithm. Mol Biol Evol, 23(10), 1891–1901.

Pond, S. L. K., Posada, D., Gravenor, M. B., Woelk, C. H., and Frost, S. D. W. (2006b). GARD: a genetic algorithm for recombination detection. Bioinformatics, 22(24), 3096–3098.

Posada, D. and Crandall, K. (1998). MODELTEST: testing the model of DNA substitution. Bioinformatics, 14(9), 817–818.

Shriner, D., Nickle, D. C., Jensen, M. A., and Mullins, J. I. (2003). Potential impact of recombination on sitewise approaches for detecting positive natural selection. Genet Res, 81(2), 115–121.

Stamatakis, A. (2014). RAxML version 8: a tool for phylogenetic analysis and post-analysis of large phylogenies. Bioinformatics, 30(9), 1312–1313.

Stern, A., Doron-Faigenboim, A., Erez, E., Martz, E., Bacharach, E., and Pupko, T. (2007). Selecton 2007: advanced models for detecting positive and purifying selection using a Bayesian inference approach. Nucleic Acids Res, 35(Web Server issue), W506–W511.

Su, F., Ou, H.-Y., Tao, F., Tang, H., and Xu, P. (2013). PSP: rapid identification of orthologous coding genes under positive selection across multiple closely related prokaryotic genomes. BMC Genomics, 14, 924.

Swanson, W. J., Nielsen, R., and Yang, Q. (2003). Pervasive adaptive evolution in mammalian fertilization proteins. Molecular biology and evolution, 20(1), 18–20.

Vitti, J. J., Grossman, S. R., and Sabeti, P. C. (2013). Detecting natural selection in genomic data. Annu Rev Genet, 47, 97–120.

Webb, A. E., Walsh, T. A., and O’Connell, M. J. (2017). VESPA: very large-scale evolutionary and selective pressure analyses. PeerJ Computer Science, 3, e118.

Yang, Z. (2007). PAML 4: phylogenetic analysis by maximum likelihood. Mol Biol Evol, 24(8), 1586–1591.

Yang, Z., Wong, W. S. W., and Nielsen, R. (2005). Bayes Empirical Bayes Inference of Amino Acid Sites Under Positive Selection. Mol Biol Evol, 22(4), 1107–1118.

